# Assessing the potential of eDNA analysis for spawning surveys in marine environments: A case study on Japanese jack mackerel

**DOI:** 10.1101/2023.09.10.557099

**Authors:** Satsuki Tsuji, Hiroaki Murakami, Reiji Masuda

**Author notes:** **e-mail addresses:** S.T.; H.M.; R.M.

## Abstract

Understanding spawning ecology is important for species/population management and conservation. However, conventional surveys are often time- and labour-consuming and invasive. To address these challenges, environmental DNA (eDNA) analysis has emerged as a promising method for spawning surveys. This study investigated the spawning season and diurnal changes in eDNA concentrations targeted Japanese jack mackerel in Maizuru Bay, Japan, to assess the potential of eDNA-based spawning surveys in the marine environment. First, to estimate the spawning season, eDNA concentrations at sunset and sunrise were compared monthly for one year at three sites. As Japanese jack mackerel are nocturnal spawners, eDNA concentration will increase at night during the spawning season. A significant increase in eDNA concentration was observed between sunset and sunrise in July, suggesting that this period corresponded to the spawning season. This was confirmed by egg collection surveys using a plankton net. Second, to investigate diurnal changes in eDNA concentrations during the spawning season, time-series sampling was conducted every three hours for 24 hours in July. The results indicated that eDNA concentration showed a significant diurnal change with a peak between 9 PM and midnight, which is suggested to be their spawning time window. This study successfully estimated the spawning season of Japanese jack mackerel and demonstrated the potential of eDNA-based spawning surveys for the first time in marine environments. These surveys are expected to contribute to the advancement of this field as a new approach to overcoming traditional challenges in marine spawning surveys.

**Highlights:**

- The potential of eDNA for spawning surveys was assessed in jack mackerel.
- Monthly morning and evening eDNA sampling identified spawning activity in July.
- Egg surveys using a plankton net confirmed the spawning event.
- Time-series sampling showed a diurnal change in eDNA concentrations in July.
- eDNA analysis is thus applicable for detecting spawning in marine species.

## Introduction

Reproductive events are an important factor in the recruitment process of fish populations because the success and magnitude of spawning directly influence population dynamics and future stock size (Mertz and Myers, 1994; Scott et al., 2006). Hence, comprehending the timing and location of fish spawning has long been recognised as a critical challenge for effective species and/or population management and conservation (Danylchuk et al., 2011; Di Muri et al., 2022; Spear et al., 2015). Conventional methods employed by fishermen, researchers, and conservation managers to identify fish spawning involve collecting fish eggs, observing the gonads of parent fish or conducting acoustic telemetry surveys. However, these methods are generally time-consuming, labour-intensive and susceptible to monitoring biases and field conditions (ex. weather, time and turbidity) (Caswell et al., 2004; Ko et al., 2013; Vautier et al., 2023). Furthermore, additional mortality is inevitable with survey methods that require the collection of individuals and eggs (Engstedt et al., 2014; Lefort et al., 2015; Wei et al., 2009). Consequently, an efficient and non-invasive method that addresses these conventional challenges would be welcomed as a new approach in fish spawning surveys.

In recent years, environmental DNA (eDNA) analysis has emerged as a rapidly advancing biodiversity assessment method and has entered a new phase where it can estimate ecological events such as spawning in addition to detecting the presence of the target species (Ip et al., 2023; Sato et al., 2021; Tsuji et al., 2022b; Wu et al., 2023). For macroorganisms, eDNA refers to DNA derived from epidermal tissue, faeces, mucus and other sources released from organisms into their habitat (Barnes and Turner, 2016; Merkes et al., 2014). In species that undergo external fertilisation, reproductive materials such as oocytes, ovarian fluid and sperm released during spawning also contribute as eDNA sources. In eDNA-based spawning surveys, the occurrence of spawning can be identified by observing rapid and significant increases in eDNA concentration and/or changes in the ratio of nuclear to mitochondrial DNA due to sperm-derived eDNA (Bylemans et al., 2017; Tsuji et al., 2022b). Since only environmental samples like water need to be collected in the field, these surveys have minimal impact on target species and their habitats, while also reducing monitoring bias linked to surveyor experience and expertise. These advantages would help overcome some of the challenges of conventional methods, leading to increasingly high expectations for eDNA analysis as a novel approach in spawning surveys.

Spawning surveys based on eDNA analysis are still in the developing stage, and their application for fish is currently limited to freshwater areas (ex. lake, Vautier et al., 2023; Wu et al., 2023; river, Bracken et al., 2019; Di Muri et al., 2022; Erickson et al., 2016; Saito et al., 2022; Tsuji and Shibata, 2025). Previous studies investigating fish which spawn in the sea, such as the Japanese jack mackerel (*Trachurus japonicus*) and the Japanese eel (*Anguilla japonica*), have reported significant increases in eDNA concentrations before and after spawning in both species when artificially spawned in laboratory conditions (Takeuchi et al., 2019; Tsuji et al., 2022b). These findings suggest that spawning surveys based on eDNA analysis could be a useful method for investigating spawning in saltwater fish. The decline of marine biodiversity is a pressing concern, with fish populations inhabiting coral reefs and seagrass beds estimated to have decreased by 34–70% in the past 40 years until 2012 due to overfishing, ecosystem destruction and climate change (WWF 2015). Considering the importance of conserving marine biodiversity and managing fisheries resources sustainably, it would be of great value to assess the feasibility of eDNA-based spawning surveys in marine environments, where water movement and currents are more complex than in freshwater.

The Japanese jack mackerel (*Trachurus japonicus*) is one of the most commercially important resources in East Asia and dominates in Maizuru Bay, Japan, from spring to autumn (Masuda, 2008; Masuda et al., 2008). While their primary spawning ground is the shelf break of the southern East China Sea (Sassa et al., 2006), small-scale regional spawning also occurs along the Japanese coast (Nishida, 2006; Xie et al., 2005). Due to the complex ocean currents surrounding the Japanese archipelago, Japanese jack mackerel spawn at different times of the year and in multiple spawning grounds, contributing to seasonal variations in stock recruitment. For example, they spawn from February to May in the main spawning ground of the East China Sea and from November to the following August in the coastal waters of Japan, with spawning occurring later in the eastern regions (Nishida, 2006). While the spawning of Japanese jack mackerel in Maizuru Bay has not been demonstrated, the otolith microstructure analysis of juveniles caught in Kunda Bay, ca. 8 km west of Maizuru Bay, suggests the presence of a population that may have been spawned near Maizuru Bay and the southern Sea of Japan (Kanaji et al., 2009). Understanding the spawning season and locations of Japanese jack mackerel in Maizuru Bay is essential to comprehending population dynamics and effectively managing resources.

This study investigated the spawning season and diurnal changes in eDNA concentrations using Japanese jack mackerel in Maizuru Bay as a model species, with the aim of assessing the applicability of eDNA-based spawning surveys in the marine environment and gathering basic information. Since changes in eDNA concentration have been suggested to be a more sensitive indicator than nuclear/mitochondrial DNA ratios in field-based eDNA spawning surveys (Saito et al., 2022; Tsuji et al., 2022b), this study used changes in the concentration of the nuclear ITS region, which is more abundant in environmental water, as a spawning indicator (Minamoto et al., 2017). In night-spawning Japanese jack mackerel, eDNA concentrations are expected to be higher before sunrise than before sunset during the spawning season. Therefore, in Experiment 1 (Exp. 1), eDNA concentrations in Japanese jack mackerel were investigated and compared once a month, before sunset and at sunrise the following day at multiple sites to estimate the spawning season. Fish eggs were then collected using plankton nets to confirm the presence or absence of spawning during the spawning season as estimated by eDNA. In Experiment 2 (Exp. 2), a time series of water samples was taken during the estimated spawning period to investigate daily changes in eDNA concentrations. Based on these results, the potential of eDNA analysis is discussed as a novel approach for conducting marine spawning surveys.

## Materials and Methods

The overview of sampling designs is presented in Fig.1. All experiments were conducted in Maizuru Bay, Japan, which has a surface area of ca. 11 km^2^ (35.481°N, 135. 332°E). The tidal range in Maizuru Bay is relatively small, remaining around 30 cm throughout the year (Japan Meteorological Agency; https://www.data.jma.go.jp/kaiyou/db/tide/suisan/suisan.php?stn=MZ).

**Figure 1.**
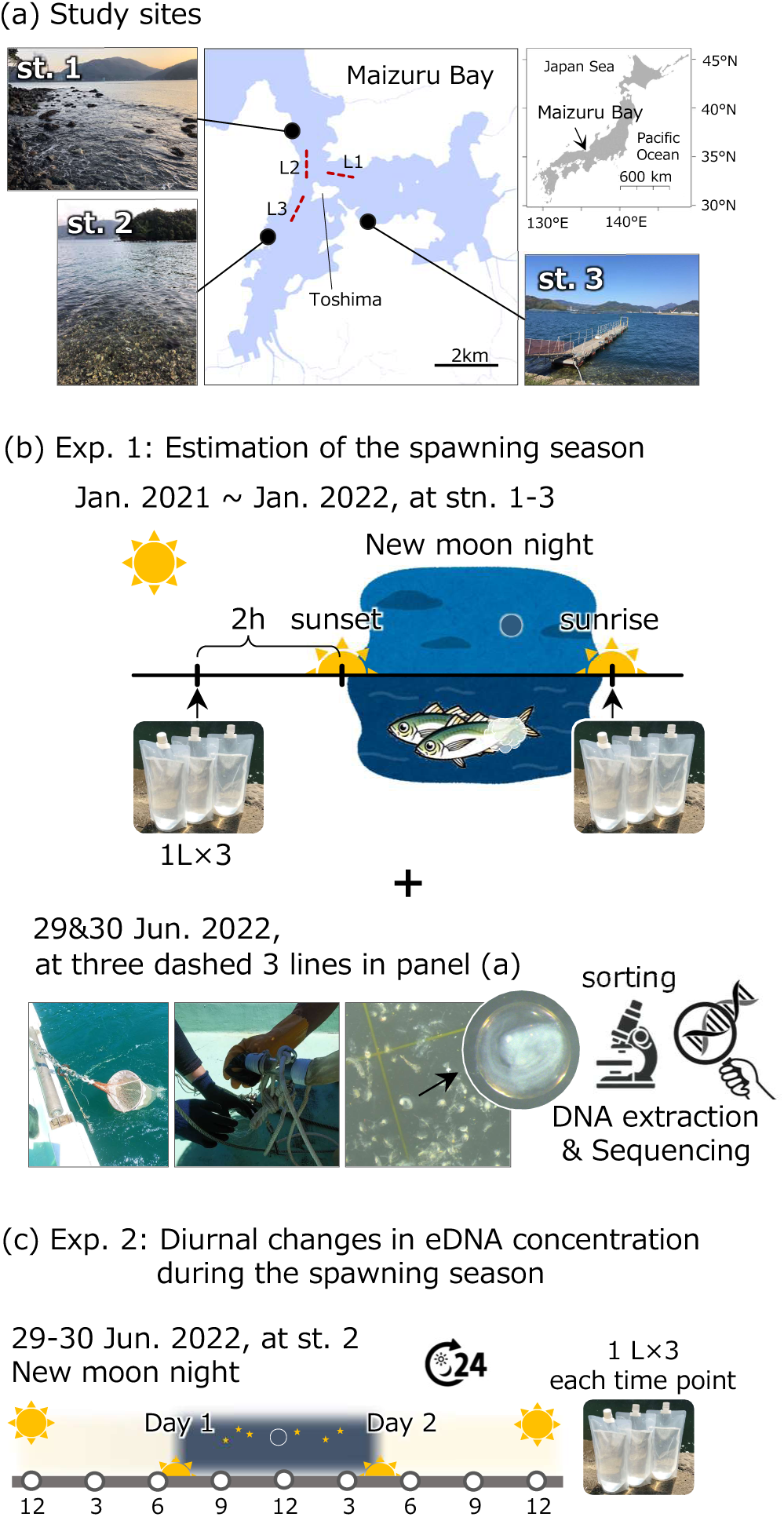
Overview of the sampling designs. (a) Study sites in Maizuru Bay, Japan, (b) and (c) sampling design for each experiment. The water sampling and the egg surveys in Exp. 1 were performed at st. 1 to 3 and L1 to 3, respectively. Time series sampling over 24 hours and every 3 hours in Exp. 2 was carried out in st. 2

### Experiment 1: Estimation of the spawning season

The water sampling survey was conducted in three sites in Maizuru Bay once a month from January 2021 to January of the following year, on the day of the new moon (Fig. 1, Table S1). We chose a new moon day on the assumption that a darker environment would be less likely to be attacked by natural enemies and spawning would be more likely to occur. As Japanese jack mackerel spawn at night; if spawning occurs, DNA concentrations are higher at sunrise than before sunset (Tsuji et al., 2022b). In each site, 1 L of surface water was collected in triplicate at two-time points, two hours before sunset and sunrise on the following day (Fig. 1b, Table S1). As Maizuru Bay is surrounded by mountains and some areas are dark before sunset, the water sampling time was set at two hours before sunset. The three duplicated samples were treated as independent samples. To preserve eDNA in water samples, benzalkonium chloride (BAC) was added and agitated well (final concentration, 0.01%; OSVAN S 10 w/v % benzalkonium chloride, Nihon Pharmaceutical). Collected water samples were promptly transported to the laboratory and vacuum-filtered using the Millipore Express PLUS disc PES (Polyethersulfone) philic filters (1 L per filter, pore size 0.45 µm, 47 mm; Merck Millipore) and filter holders (PP-47; ADVANTEC) within one hour of collection. To avoid contamination, filter holders for the number of samples were decontaminated by prior soaking in 10% hypochlorite for at least two hours. After filtration of the sunrise samples, 1 L of ultrapure water was filtered in the same way as the collected water samples and used as a filter negative control (Filt-NC). All filter samples were folded in half, wrapped in aluminium foil and immediately frozen at −20°C. DNA was extracted from the filter samples, and the nuclear internal transcribed spacer-1 region (ITS1) of the ribosomal RNA gene of Japanese jack mackerel was quantified using quantitative real-time PCR and species-specific primer-probe sets (detailed below).

To confirm the presence or absence of spawning during the spawning season (July; see Results) as estimated by eDNA, egg surveys were conducted on 29 (9:10–10:17 AM) and 30 (1:09–1:58 PM) June 2022 at three sites in Maizuru Bay (dashed lines, L1 to L3, in Fig. 1a). It was preferred to conduct egg surveys on the new moon of July when spawning was shown to be possible, but in 2022 the new moon was on 28-29 July. Thus, egg surveys were conducted at the end of June, i.e. at a new moon closer to 10 July 2021. Oblique tawing of a plankton net (diameter 450 mm, mesh 0.3 mm) was performed three or four times for each site (average 3 min 10 sec trawl per trial, wire length 40 m, wire angle 63–78°). Tables S3 and S4 show the towing conditions for each trawling and water quality at each site. The collections were pooled for each site and immediately stored on ice. In the laboratory, approximately 0.6–1.2 mm sized spherical fish eggs were sorted under a stereomicroscope and then fixed with 70% ethanol. For each collected egg, DNA was extracted using the spin column (EconoSpin, EP-31201; GeneDesign Inc.) and the buffer reagents supplied in the DNeasy Blood and Tissue kit (Qiagen). Eggs were soaked in a mixture of 180 µL buffer ATL and 20 µL Proteinase K and incubated at 56°C for 3 hours to dissolve completely. 200 µL of ethanol was added and mixed well, after which the entire elute was transferred to a spin column. After centrifugation at 6,000 g for 1 min, DNA was purified according to the manufacturer’s protocol of the DNeasy Blood and Tissue kit. Finally, the DNA was eluted in 50 µL of Buffer AE and stored at −20°C until use in the PCR assay. The species identification was conducted based on the sequencing of the mitochondrial 12S rRNA region with the MiFish fish-universal primer set (detailed below). Based on current laws and guidelines of Japan relating to animal experiments on fish, the collection of fish eggs and the use of DNA samples are allowed without any ethical approvals from any authorities.

### Experiment 2: Observation of diurnal changes in eDNA concentration during the spawning season

Time-series water sampling was conducted between 29 and 30 June 2022. At site 2, where the most significant increase in eDNA concentration was observed between sunset and sunrise in Exp. 1 (see Results), 1 L of surface water was collected in triplicate at nine time points: 12:00 noon, 3:00, 6:00, 9:00 PM, 12:00 midnight, 3:00, 6:00, 9:00 AM, and 12:00 noon (Table S2). Each water sample was added with BAC (final concentration, 0.01%) and filtered on-site using the same techniques and equipment as in Experiment 1. After filtration of 12:00 samples on 30 June, 1 L of ultrapure water was filtered as a filter negative control (Filt-NC). All filter samples were folded in half, wrapped in aluminium foil and immediately frozen at −20°C. DNA was extracted from the filter samples, and ITS1 region of Japanese jack mackerel was quantified using the same way as in Experiment 1.

### DNA extraction from filter samples

A PES filter sample was folded in quarters and placed in the lower part of the spin column (EconoSpin) with the silica gel membrane preliminarily removed. A mixed extraction buffer comprising 200 µL ultrapure water, 200 µL Buffer AL (Qiagen), and 20 µL Proteinase K was added to each filter, and the spin columns were incubated for 30 min at 56°C. The filter was moved to the top of the spin column and then centrifuged at 6,000 g for 1 min. 600 μL of ethanol was added to the collected liquid and mixed well by pipetting. The DNA in the mixture was purified using a DNeasy Blood and Tissue kit following the manufacturer’s protocol. Finally, the DNA was eluted in 100 µL of Buffer AE and stored at −20°C until use in the quantitative real-time PCR assay.

### Quantitative real-time PCR (qPCR) assays

Quantitative real-time PCRs were performed in triplicate using the Light Cycler 96 system (Roche, Basel, Switzerland). The ITS regions of Japanese jack mackerel were amplified with species-specific primer-probe sets, which were developed by previous studies (Jo et al., 2019). The sequences of the primers and a probe were as follows: Tja_ITS1_F, 5′-GCG GGT ACC CAA CTC TCT TC-3′; Tja_ITS1_R, 5′-CCT GAG CGG CAC ATG AGA G-3′ and Tja_ITS1_Pr, 5′-[FAM]-CTC TCG CTT CTC CGA CCC CGG TCG-[BHQ1]-3′. The qPCR was conducted in a total volume of 15 μL, containing 900 nM of forward and reverse primers, 125 nM TaqMan probe, 0.075 μL of AmpErase uracil N-glycosylase (Thermo Fisher Scientific), 7.5 μL of 2 × TaqMan Environmental Master Mix 2.0 (Thermo Fisher Scientific), and 2 μL of DNA template. A four-step dilution series containing 3 × 10^1^ to 3 × 10^4^ copies was used for each qPCR run as quantification standards. As PCR-negative controls, three reactions were included in all qPCR runs in which 2.0 μL of ultrapure water was added instead of the DNA template. The qPCR thermal conditions were as follows: 2 min at 50°C, 10 min at 95°C, then 55 cycles of 15 s at 95°C, and 75 s at 60°C. To avoid contamination, reagent preparation and qPCR were carried out in separate rooms. The R^2^ values for the standard curve of all qPCR were shown in Table S5. No amplification was observed in any of the Filt-NC and PCR-negative controls.

### Species identification of eggs

To identify fish species of each egg, approximately 170 base pair of 12S rRNA region was amplified by a two-step tailed PCR approach with two MiFish fish universal primer sets (Table S6, Miya et al., 2015). The first-round PCR (1stPCR) was performed in 12-µL reactions containing 6 µL of 2×KODone PCR Master Mix (TOYOBO), 1.4 μL of each primer mixture (MiFish-U: MiFish-U2 = 10: 3; 5µM) and 2 μl of template DNA. There was no PCR replication. The thermal conditions were as follows: 30 cycles of 10 sec at 98°C, 5 sec at 65°C and 5 sec at 68°C. Amplicons of the 1stPCR were purified using Sera-Mag SpeedBeads Carboxylate-Modified magnetic particles (Hydrophobic) (Cytiva) (PCR product : Sera-Mag beads = 1 : 0.8). The purified amplicons were adjusted to 0.1 ng/µL and used as the template in second-round PCR (2ndPCR). The 2ndPCR was performed in 12-µL reactions containing 6.0 μL of 2 × KAPA HiFi HotStart ReadyMix (KAPA Biosystems), 2.0 μL of each primer with index (1.8 μM) and 2.0 μL of the purified 1stPCR amplicon. The sequences of the 2ndPCR primers were shown in Table S6. The thermal conditions were as follows: 3 min at 95°C, 12 cycles of 20 s at 98°C, 15 s at 72°C and 5 min at 72°C. The indexed 2nd PCR products were pooled in equal volumes, and the target band (approx. 370 bp) was excised using a 2% E-Gel SizeSelect Agarose Gels (Thermo Fisher Scientific). The 150 bp paired-end sequencing by NovaSeq was outsourced to Novogene Co., Ltd. All raw sequences were deposited in the DDBJ Sequence Read Archive (accession number: DRA016903). The bioinformatics analysis was performed using the PMiFish pipeline (https://github.com/rogotoh/PMiFish). The top BLAST hit with a sequence identity ≥ 98% was identified as the species of the egg.

### Statistical analysis

All statistical analyses and graphic illustrations for this study were conducted using R version 4.2.2 software (R Core Team, 2023). In experiment 1, we performed a Wilcoxon rank sum test to compare the eDNA concentration between 2-hours before sunset and sunrise samples by wilcox.exact function in exactRankTests package ver. 0.8.34. To account for the multiplicity of tests, the Bonferroni correction was applied, and the α level of significance was set at 0.003.

In experiment 2, to examine diurnal changes in eDNA concentration, the ITS1 concentration observed during the daytime (12:00 noon, 3:00 PM, 6:00PM of 29 June 2022, 6:00 AM, 9:00 AM, 12:00 noon of 30 June 2022) was compared with those observed at each time point during the nighttime (9:00 PM, 12:00 midnight of 29 of June 2022, 3:00 AM of 30 June 2022) using the Kruskal–Wallis test followed by the Conover’s test with Holm adjustment by kwAllPairsConoverTest function in PMCMRplus package ver.1.9.3. The α level of significance was set at 0.05.

## Results

Japanese jack mackerel eDNA was detected and successfully quantified at all sites throughout the year. A comparison of eDNA concentrations at two hours before sunset (hereafter “sunset”) and sunrise in each month showed significantly higher ITS1 concentrations at sunrise than at sunset in July (*p* < 0.001; Fig. 2). A similar trend was observed in March and October, but it was not statistically significant (*p* = 0.02 and *p* = 1.0, respectively). On the other hand, in February and September, eDNA concentrations were significantly higher at sunset than at sunrise (*p* < 0.001).

**Figure 2.**
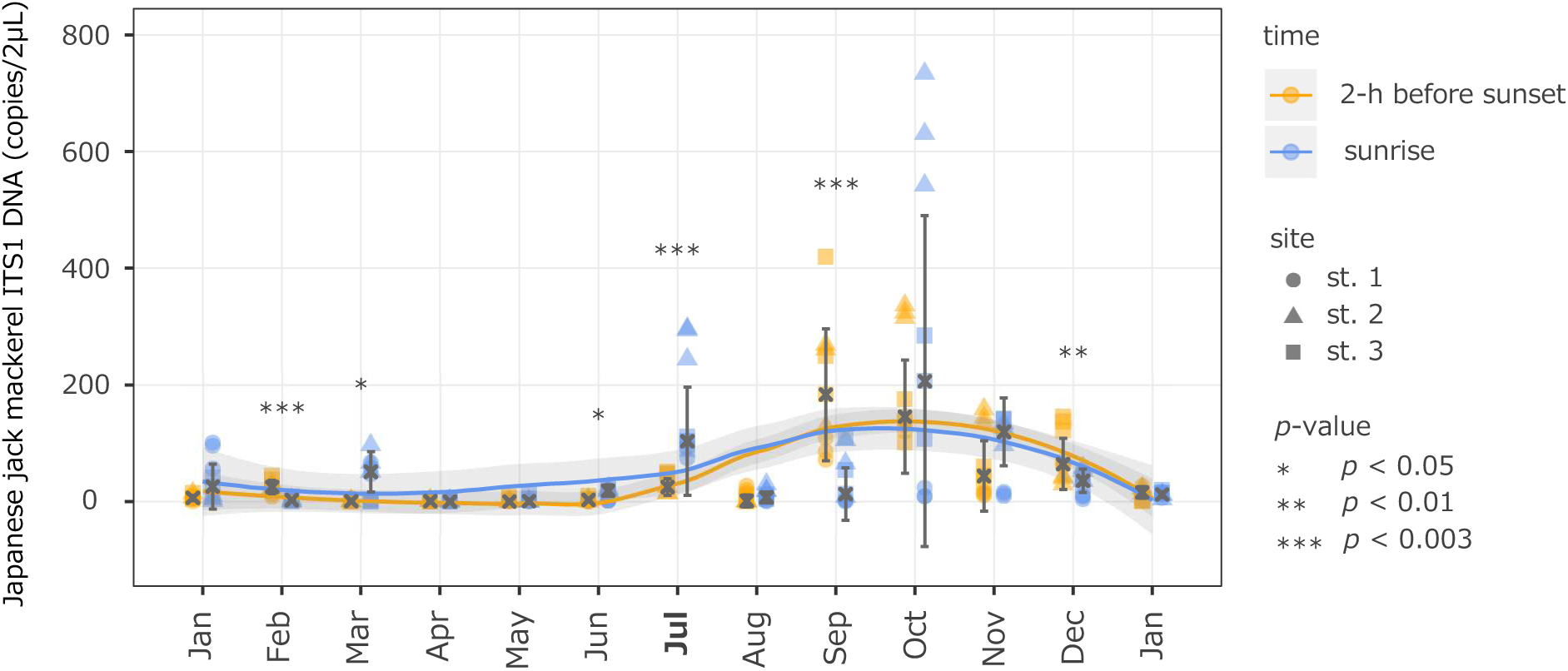
Changes in eDNA concentrations of Japanese jack mackerel two hours before sunset and at sunrise on the following day in each month. The cross mark with error bars indicates median values and standard deviations. Lines and grey areas indicate a loess regression and 95% CI. Significant differences in eDNA concentrations between two hours before sunset and at sunrise are indicated by asterisks (Wilcoxon rank sum test). To account for the multiplicity of tests, the Bonferroni correction was applied, and the α level of significance was set at 0.003.

In the egg collection survey, 260 fish eggs were successfully identified to species (or taxon) by sequencing, of which two were identified as Japanese jack mackerel eggs. These were collected from line L3 in Fig. 1a. The remaining eggs were assigned to species inhabiting Maizuru Bay (Fig. S1A). In total, 25 species were identified, representing approximately 90% of the expected species diversity based on the species accumulation curve (Fig. S1b, c, Table S7). Eggs of species with relatively low occurrence frequencies, including Japanese jack mackerel, were detected on only one of the two survey days (30 June 2022).

Diurnal changes in eDNA concentrations were observed in the time-series water sampling at site 2 (Fig. 3). Environmental DNA concentrations were significantly higher at nighttime than during the daytime (*P* < 0.01 for 9:00 PM and 12:00 midnight, *P* < 0.05 for 3:00 AM, Conover’s test), peaking between 21:00 and 24:00.

**Figure 3.**
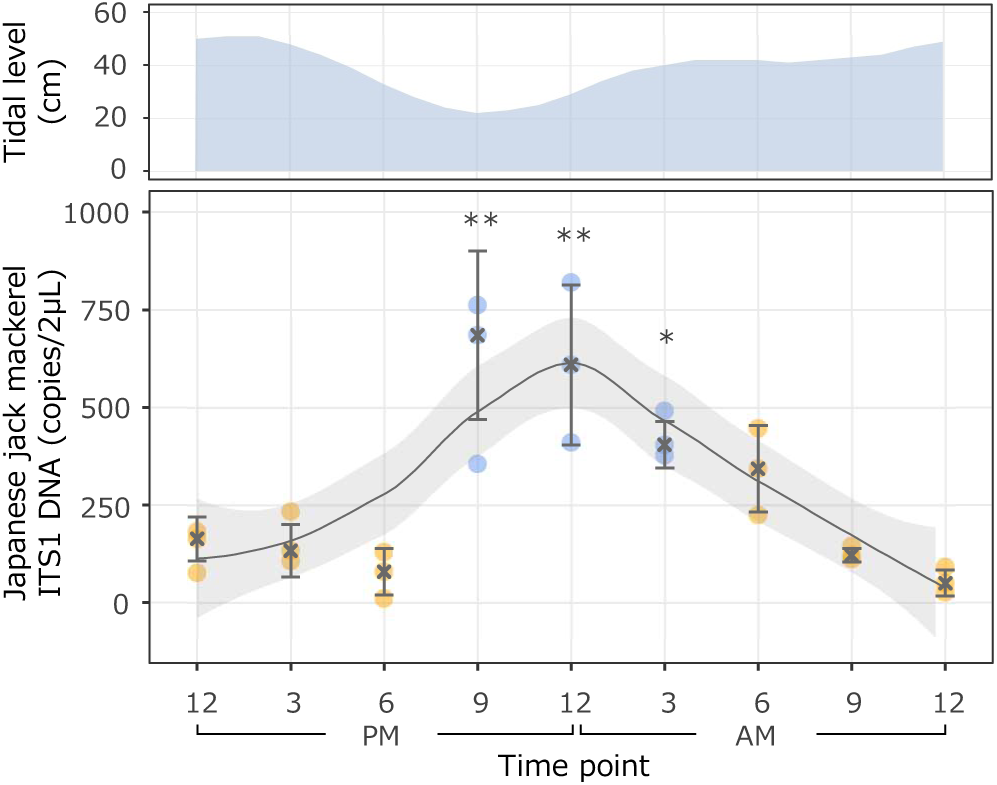
Daily changes in tidal levels in Maizuru Bay (top panel) and diurnal changes in eDNA concentrations of Japanese jack mackerel ITS1 at st. 2 (bottom panel) from 29 to 30 June 2022. The orange and blue plot indicates the daytime (12:00 noon, 3:00 PM, 6:00 PM, 6:00 AM, 9:00 AM and day 2 12:00 noon) and nighttime (9:00 PM, 12:00 midnight and 3:00 AM), respectively. Line and grey area indicate a loess regression and 95% CI. The ITS1 concentration observed during the daytime was compared with those observed at each time point during the nighttime. The *p* values are indicated by asterisks (Conover’s test; * *p* < 0.05, ** *p* < 0.01). Sunset and sunrise times are as follows: 7:18 PM and 4:46 AM.

## Discussion

In this study, eDNA analysis was successfully employed to estimate the spawning season of Japanese jack mackerel in Maizuru Bay and revealed that eDNA concentrations change diurnally during the spawning season in relation to spawning activity. The eDNA-based estimation of spawning season was supported by the capture of their eggs using trawling with a plankton net. This study is the first to demonstrate the applicability of eDNA analysis for fish spawning surveys in the marine environment and highlights its potential utility. Furthermore, understanding the dynamics of eDNA concentrations during the spawning season provides valuable insights not only for future eDNA-based spawning surveys but also for species diversity monitoring.

During monthly eDNA monitoring throughout the year, the increase in nighttime eDNA concentrations observed in July was considered to reflect the occurrence of Japanese jack mackerel spawning (Fig. 2). At site 2, the most significant increases in eDNA concentrations were observed between sunset and sunrise, suggesting that relatively more spawning occurred near there (Tsuji and Shibata, 2021). In Wakasa Bay, the outer bay of Maizuru Bay, two groups of Japanese jack mackerel with distinct morphologies and ecological characteristics coexist: the offshore, seasonally migrating group (black-type) and the coastal, non-migratory group (yellow-type). Observations of body length composition and otoliths of their juveniles caught in Wakasa Bay suggested that black-type and yellow-type are derived from eggs spawned between January and March and between June to July, respectively (Azeta and Ochiai, 1962). The presence of cohorts presumably hatched between mid-June and early August has also been observed in Kunda Bay, located to the west of Maizuru Bay (Kanaji et al., 2009). Taken together, the July spawning estimated by eDNA analysis is likely attributable to the coastal yellow-type. Furthermore, while the water depth range in the main spawning grounds has been estimated between 100 to 200 m (Nishida 2006), Maizuru Bay has a maximum depth of approximately 30 meters near the bay’s mouth and north of Toshima Island, and the depth is generally shallower towards the back of the bay (cf. Minami et al., 2018). Thus, the depths of Maizuru Bay are not quite comparable to those of the main spawning grounds, but if Japanese jack mackerel spawn in the bay, they are likely to prefer deeper sites. The tidal currents are strong from the mouth of the bay to Toshima; therefore, assuming that spawning occurs around Toshima, where the currents begin to slow, the most noticeable increase in eDNA concentration observed at site 2 appears reasonable (Miwa and Ikeno, 2008). Additionally, if spawning takes place in deeper areas, more significant changes in eDNA concentrations could have been detected by sampling deeper water rather than surface water (Fukumori et al., 2024). In this study, due to cost, number of researcher and safety limitations, it was only possible to sample surface water from the shore. Future studies may achieve more sensitive detection of spawning-induced increases in eDNA concentrations by conducting water sampling at depths corresponding to the spawning sites.

Increases in eDNA concentrations between sunset and sunrise were also observed at two out of three sites in March and October, but this could reflect migration-related noise given the ecology of Japanese jack mackerel. In March, the quantified eDNA concentrations were relatively low at all sites, both at sunset and sunrise, making it unlikely that spawning behaviour involving the release of large amounts of sperm occurred. On the other hand, October is the period when juveniles that entered from the outer bay during the summer are growing and actively swimming in the bay (Masuda et al., 2008). Therefore, relatively high eDNA concentrations were observed at sunset at all sites. Although there was an increase in concentrations at sites 2 and 3 on sunrise, the increase was only 1.4 to 1.9-fold, which is not remarkable increase. The amount of faeces, one of the sources of eDNA, is likely to increase as fish actively feed in the early morning. Therefore, it is reasonable to assume that the changes in eDNA concentrations observed in October were most likely caused by fish migration and increased activity. Additionally, the significantly higher DNA concentrations before sunset than at sunrise, observed in February and September, were also considered to be due to fish migration and feeding, as Japanese jack mackerel spawn at night.

The occurrence of Japanese jack mackerel spawning in Maizuru Bay during July (the end of June), as estimated based on eDNA analysis, was confirmed by the successful direct recovery of their eggs. Although the fish egg recovery surveys conducted in this study were not of an experimental design that provided quantitative data, given that the Japanese jack mackerel is one of the dominant species in Maizuru Bay, the fact that only two eggs were recovered likely indicates a low spawning quantity. This is consistent with our assumption that the yellow-type, which has a smaller biomass than the black-type, is spawning (see above). Based on the water temperature during the survey (20–25°C, Table SX), the recovered Japanese jack mackerel eggs may have been floating for a maximum of one day (Ochiai et al., 1982). Therefore, the possibility that the eggs were introduced from outside the bay cannot be completely ruled out, depending on the tidal current conditions. However, as it would have taken more than a day for the eggs to flow from the outer bay to Maizuru Bay, we concluded that the eggs recovered in this study were likely spawned within the bay or at least nearshore waters close to the bay mouth.

The significant diurnal changes in eDNA concentrations observed in the time-series sampling suggested that Japanese jack mackerel mainly spawn between 9:00 PM and 12:00 midnight (Fig. 3). The possibility that eDNA concentrations were influenced by tidal effects was ruled out due to the lack of agreement in their respective patterns of change. The spawning time window of Japanese jack mackerel under natural conditions has been predicted to be before dawn, but this has not been definitively identified and remained unknown (Nishida, 2006; Yoda et al., 2006). However, our results are consistent with the peak spawning window observed in previous studies where they were artificially spawned in tanks (Nyuji et al., 2013). It is also consistent with information on other *Trachurus* species, *T. symmetricus* and *T. trachurus*, which spawn at night (Karlou-Riga and Economidis, 1997; Macewicz and Hunter, 1993). These facts would emphasise the need for a re-investigation of the spawning time window of Japanese jack mackerel under natural conditions. Diurnal changes in eDNA concentrations during the spawning season have been reported in only a single study, which focuses on *Plecoglossus altivelis* in riverine systems (Tsuji and Shibata, 2025). Our study extends this understanding by revealing that such diurnal changes, associated with spawning activities, also occur in marine environments where water flow is more complex and multidirectional. A sampling plan aligned with the spawning time window of the target species is likely to maximise the sensitivity of eDNA-based spawning surveys (Inui et al., 2021). Furthermore, since eDNA concentrations are often used as indicators of species biomass (Rourke et al., 2022), considering changes in DNA concentrations during the spawning season in general eDNA-based biological monitoring could improve the reliability of the results. Information on the dynamics of eDNA related to spawning behaviour is very limited, despite its potential to be effectively used in various aspects of studies using eDNA analysis, from sampling planning to result interpretation (Tsuji et al., 2022b; Wu et al., 2022). Future studies should carefully investigate the diffusion, advection, degradation, and interspecific differences of eDNA etc. associated with spawning.

In conclusion, this study has demonstrated the potential utility of eDNA-based spawning surveys in the marine environment by successfully estimating the spawning season of Japanese jack mackerel in Maizuru Bay. Furthermore, the study revealed that eDNA concentrations show diurnal changes during the spawning season with a peak at the time window of spawning, providing useful information not only for spawning surveys using eDNA analysis but also for species diversity surveys in general. While only Japanese jack mackerel was focused on in this study, future research could potentially estimate the spawning ecology of multiple species within the target taxonomic group simultaneously by employing quantitative eDNA metabarcoding methods (Tsuji et al., 2022a; Ushio et al., 2018). Currently, eDNA analysis cannot provide some information obtained through conventional methods, such as fish size, age, sex, and hybridisation (Minamoto, 2022); however, it has the advantage of efficiently and non-invasively estimating spawning seasons. This characteristic could significantly alleviate the challenges associated with spawning surveys in marine environments and contribute to advancements in this research field.

## Acknowledgements

We wish to thank Dr. Keita W. Suzuki, Ms. Misaki Shiomi, Mr. Hunter Godfrey and Mr. Yoshihito Ogura (Maizuru Fisheries Research Station, Kyoto University) for their help with the collection of fish eggs in experiment 1. This study was supported by JSPS KAKENHI Grant Number JP20K15578 and JP19H05641.

## Conflict of Interest Statement

The authors have no conflicts of interest to declare.

## Authors’ contributions

S.T.: Conceptualization, Funding acquisition, Methodology, Fieldwork (all experiments), Molecular analysis, Visualization, Writing−original draft. H.M.: Methodology, Fieldwork (experiment 1), Writing−review & editing. R.M.: Conceptualization, Funding acquisition, Methodology, Fieldwork (experiment 1), Writing−review & editing, Supervision.

## Data availability

Full details of the qPCR results for each experiment of the present study are available in the supporting information (Table S8 and S9). All raw sequences were deposited in the DDBJ Sequence Read Archive (accession number: DRA016903).

---

Figure S1. (a) species and their frequency of the collected eggs by egg surveys, (b) the number of detected species per survey date; (c) the number of analysed eggs and species diversity.

## Notes

### Competing Interest Statement

The authors have declared no competing interest.

### Summary of Updates

The title has been changed to match the content of the manuscript. The introduction and discussion have been significantly revised to focus more on the applicability of environmental DNA analysis to spawning surveys in marine environments. Some figures have also been merged and the overall number of figures reduced. Table 1 has been deleted.

## References

Azeta M., Ochiai A., 1962. A study on the race of Jack mackerel found in Wakasa Bay. NIPPON SUISAN GAKKAISHI 28, 967–978. 10.2331/suisan.28.967

Barnes, M.A., Turner, C.R., 2016. The ecology of environmental DNA and implications for conservation genetics. Conservation Genetics 17, 1–17. 10.1007/s10592-015-0775-4

Bracken, F.S.A., Rooney, S.M., Kelly-Quinn, M., King, J.J., Carlsson, J., 2019. Identifying spawning sites and other critical habitat in lotic systems using eDNA “snapshots”: A case study using the sea lamprey *Petromyzon marinus* L. Ecology and Evolution 9, 553–567. 10.1002/ece3.4777

Bylemans, J., Furlan, E.M., Hardy, C.M., McGuffie, P., Lintermans, M., Gleeson, D.M., 2017. An environmental DNA-based method for monitoring spawning activity: A case study, using the endangered Macquarie perch (*Macquaria australasica*). Methods in Ecology and Evolution 8, 646–655. 10.1111/2041-210X.12709

Caswell, N.M., Peterson, D.L., Manny, B.A., Kennedy, G.W., 2004. Spawning by lake sturgeon (*Acipenser fulvescens*) in the Detroit River. Journal of Applied Ichthyology 20, 1–6. 10.1111/j.1439-0426.2004.00499.x

Danylchuk, A.J., Cooke, S.J., Goldberg, T.L., Suski, C.D., Murchie, K.J., Danylchuk, S.E., Shultz, A.D., Haak, C.R., Brooks, E.J., Oronti, A., Koppelman, J.B., Philipp, D.P., 2011. Aggregations and offshore movements as indicators of spawning activity of bonefish (*Albula vulpes*) in The Bahamas. Marine Biology 158, 1981–1999. 10.1007/s00227-011-1707-6

Di Muri, C., Lawson Handley, L., Bean, C.W., Benucci, M., Harper, L.R., James, B., Li, J., Winfield, I.J., Hänfling, B., 2023. Spatio-temporal monitoring of lake fish spawning activity using environmental DNA metabarcoding. Environmental DNA 5, 849–860.. 10.1002/edn3.343

Engstedt, O., Engkvist, R., Larsson, P., 2014. Elemental fingerprinting in otoliths reveals natal homing of anadromous Baltic Sea pike (*Esox lucius* L.). Ecology of Freshwater Fish 23, 313–321. 10.1111/eff.12082

Erickson, R.A., Rees, C.B., Coulter, A.A., Merkes, C.M., McCalla, S.G., Touzinsky, K.F., Walleser, L., Goforth, R.R., Amberg, J.J., 2016. Detecting the movement and spawning activity of bigheaded carps with environmental DNA. Molecular Ecology Resources 16, 957–965. 10.1111/1755-0998.12533

Fukumori, K., Kondo, N.I., Kohzu, A., Tsuchiya, K., Ito, H., Kadoya, T., 2024. Vertical eDNA distribution of cold-water fishes in response to environmental variables in stratified lake. Ecology and Evolution 14, e11091. 10.1002/ece3.11091

Inui, R., Akamatsu, Y., Kono, T., Saito, M., Miyazono, S., Nakao, R., 2021. Spatiotemporal changes of the environmental DNA concentrations of amphidromous fish *Plecoglossus altivelis altivelis* in the spawning grounds in the Takatsu River, western Japan. Frontiers in Ecology and Evolution 9, 182. 10.3389/fevo.2021.622149

Ip, Y.C.A., Chang, J.J.M., Tun, K.P.P., Meier, R., Huang, D., 2023. Multispecies environmental DNA metabarcoding sheds light on annual coral spawning events. Molecular Ecology 32, 6474–6488. 10.1111/mec.16621

Jo, T., Arimoto, M., Murakami, H., Masuda, R., Minamoto, T., 2019. Particle size distribution of environmental DNA from the nuclei of marine fish. Environmental Science & Technology 53, 9947–9956. 10.1021/acs.est.9b02833

Kanaji, Y., Watanabe, Y., Kawamura, T., Xie, S., Yamashita, Y., Sassa, C., Tsukamoto, Y., 2009. Multiple cohorts of juvenile jack mackerel *Trachurus japonicus* in waters along the Tsushima Warm Current. Fisheries Research 95, 139–145. 10.1016/j.fishres.2008.08.004

Karlou-Riga, C., Economidis, P.S., 1997. Spawning frequency and batch fecundity of horse mackerel, *Trachurus trachurus* (L.), in the Saronikos Gulf (Greece). Journal of Applied Ichthyology 13, 97–104. 10.1111/j.1439-0426.1997.tb00108.x

Ko, H.-L., Wang, Y.-T., Chiu, T.-S., Lee, M.-A., Leu, M.-Y., Chang, K.-Z., Chen, W.-Y., Shao, K.-T., 2013. Evaluating the accuracy of morphological identification of larval fishes by applying DNA barcoding. PLOS One 8, e53451. 10.1371/journal.pone.0053451

Lefort, M.-C., Boyer, S., Barun, A., Khoyi, A.E., Ridden, J., Smith, V.R., Sprague, R., Waterhouse, B.R., Cruickshank, R.H., 2015. Blood, sweat and tears: non-invasive vs. non-disruptive DNA sampling for experimental biology (No. e1580). PeerJ Inc. 10.7287/peerj.preprints.655v3

Macewicz, B.J., Hunter, R.J., 1993. Spawning frequency and batch fecundity of jack mackerel, *Trachurus symmetricus*, off California during 1991. Calif. Coop. Oceanic Fish. Invest. Rep. 34, 112–121.

Masuda, R., 2008. Seasonal and interannual variation of subtidal fish assemblages in Wakasa Bay with reference to the warming trend in the Sea of Japan. Environmental Biology of Fishes 82, 387–399.

Masuda, R., Yamashita, Y., Matsuyama, M., 2008. Jack mackerel *Trachurus japonicus* juveniles use jellyfish for predator avoidance and as a prey collector. Fisheries Science 74, 276–284.

Merkes, C.M., McCalla, S.G., Jensen, N.R., Gaikowski, M.P., Amberg, J.J., 2014. Persistence of DNA in carcasses, slime and avian feces may affect interpretation of environmental DNA data. PLOS One 9, e113346.

Mertz, G., Myers, R.A., 1994. Match/mismatch predictions of spawning duration versus recruitment variability. Fisheries Oceanography 3, 236–245. 10.1111/j.1365-2419.1994.tb00101.x

Minami, K., Sawada, H., Masuda, R., Takahashi, K., Shirakawa, H., Yamashita, Y., 2018. Stage-specific distribution of Japanese sea cucumber *Apostichopus japonicus* in Maizuru Bay, Sea of Japan, in relation to environmental factors. Fisheries Science 84, 251–259. 10.1007/s12562-017-1174-1

Minamoto, T., 2022. Environmental DNA analysis for macro-organisms: species distribution and more. DNA Research 29, dsac018. 10.1093/dnares/dsac018

Minamoto, T., Uchii, K., Takahara, T., Kitayoshi, T., Tsuji, S., Yamanaka, H., Doi, H., 2017. Nuclear internal transcribed spacer-1 as a sensitive genetic marker for environmental DNA studies in common carp *Cyprinus carpio*. Molecular Ecology Resources 17, 324–333. 10.1111/1755-0998.12586

Miwa H., Ikeno H., 2008. Numerical analysis of tidal current and water quality in stratified field on summer in Maizuru Bay. Annual Journal of Hydraulic Engineering 52, 1387–1392. 10.2208/prohe.52.1387

Miya, M., Sato, Y., Fukunaga, T., Sado, T., Poulsen, J.Y., Sato, K., Minamoto, T., Yamamoto, S., Yamanaka, H., Araki, H., 2015. MiFish, a set of universal PCR primers for metabarcoding environmental DNA from fishes: detection of more than 230 subtropical marine species. Royal Society Open Science 2, 150088.

Nishida H., 2006. Reproductive Biology of Japanese Jack Mackerel *Trachurus japonicus* and Japanese Sardine *Sardinops melanostictus*. Bulletin of Fisheries Research Agency. Supplement 113–118.

Nyuji, M., Fujisawa, K., Imanaga, Y., Kitano, H., Yamaguchi, A., Matsuyama, M., 2013. GnRHa-induced spawning of wild-caught jack mackerel *Trachurus japonicus*. Fisheries Science 79, 251–258. 10.1007/s12562-013-0599-4

Ochiai, A., Mutsutani, K., Umeda, S., 1982. Development of eggs, larvae and juveniles of jack mackerel, *Trachurus japonicus*. Japanese Journal of Ichthyology 29, 86–92. 10.11369/jji1950.29.86

R Core Team, 2023. R: A language and environment for statistical computing. R Foundation for Statistical Computing.

Rourke, M.L., Fowler, A.M., Hughes, J.M., Broadhurst, M.K., DiBattista, J.D., Fielder, S., Wilkes Walburn, J., Furlan, E.M., 2022. Environmental DNA (eDNA) as a tool for assessing fish biomass: A review of approaches and future considerations for resource surveys. Environmental DNA 4, 9–33. 10.1002/edn3.185

Saito, M., Tsuji, S., Nakao, R., Miyazono, S., Akamatsu, Y., 2022. Comparative study on nuclear and mitochondrial DNA of Ayu *Plecoglossus altivelis* for environmental DNA-based spawning evaluation. Landscape and Ecological Engineering 19, 55–67. 10.1007/s11355-022-00519-5

Sassa, C., Konishi, Y., Mori, K., 2006. Distribution of jack mackerel (*Trachurus japonicus*) larvae and juveniles in the East China Sea, with special reference to the larval transport by the Kuroshio Current. Fisheries Oceanography 15, 508–518. 10.1111/j.1365-2419.2006.00417.x

Sato, M., Inoue, N., Nambu, R., Furuichi, N., Imaizumi, T., Ushio, M., 2021. Quantitative assessment of multiple fish species around artificial reefs combining environmental DNA metabarcoding and acoustic survey. Scientific Reports 11, 1–14.

Scott, B.E., Marteinsdottir, G., Begg, G.A., Wright, P.J., Kjesbu, O.S., 2006. Effects of population size/age structure, condition and temporal dynamics of spawning on reproductive output in Atlantic cod (*Gadus morhua*). Ecological Modelling 191, 383–415. 10.1016/j.ecolmodel.2005.05.015

Spear, S.F., Groves, J.D., Williams, L.A., Waits, L.P., 2015. Using environmental DNA methods to improve detectability in a hellbender (*Cryptobranchus alleganiensis*) monitoring program. Biological Conservation 183, 38–45. 10.1016/j.biocon.2014.11.016

Takeuchi, A., Iijima, T., Kakuzen, W., Watanabe, S., Yamada, Y., Okamura, A., Horie, N., Mikawa, N., Miller, M.J., Kojima, T., 2019. Release of eDNA by different life history stages and during spawning activities of laboratory-reared Japanese eels for interpretation of oceanic survey data. Scientific Reports 9, 1–9.

Tsuji, S., Inui, R., Nakao, R., Miyazono, S., Saito, M., Kono, T., Akamatsu, Y., 2022a. Quantitative environmental DNA metabarcoding shows high potential as a novel approach to quantitatively assess fish community. Scientific Reports 12, 21524. 10.1038/s41598-022-25274-3

Tsuji, S., Murakami, H., Masuda, R., 2022b. Analysis of the persistence and particle size distributional shift of sperm-derived environmental DNA to monitor jack mackerel spawning activity. Environmental Science & Technology 56, 10754–10763. 10.1021/acs.est.2c01904

Tsuji, S., Shibata, N., 2025. Particle size distribution shift and diurnal concentration changes of environmental DNA caused by fish spawning behaviour. Landscape and Ecological Engineering 21, 151–161. 10.1007/s11355-024-00630-9

Tsuji, S., Shibata, N., 2021. Identifying spawning events in fish by observing a spike in environmental DNA concentration after spawning. Environmental DNA 3, 190–199. 10.1002/edn3.153

Ushio, M., Murakami, H., Masuda, R., Sado, T., Miya, M., Sakurai, S., Yamanaka, H., Minamoto, T., Kondoh, M., 2018. Quantitative monitoring of multispecies fish environmental DNA using high-throughput sequencing. Metabarcoding and Metagenomics 2, e23297. 10.3897/mbmg.2.23297

Vautier, M., Chardon, C., Goulon, C., Guillard, J., Domaizon, I., 2023. A quantitative eDNA-based approach to monitor fish spawning in lakes: Application to European perch and whitefish. Fisheries Research 264, 106708. 10.1016/j.fishres.2023.106708

Wei, Q.W., Kynard, B., Yang, D.G., Chen, X.H., Du, H., Shen, L., Zhang, H., 2009. Using drift nets to capture early life stages and monitor spawning of the Yangtze River Chinese sturgeon (*Acipenser sinensis*). Journal of Applied Ichthyology 25, 100–106. 10.1111/j.1439-0426.2009.01269.x

Wu, L., Wu, Q., Inagawa, T., Okitsu, J., Sakamoto, S., Minamoto, T., 2023. Estimating the spawning activity of fish species using nuclear and mitochondrial environmental DNA concentrations and their ratios. Freshwater Biology 68, 103–114. 10.1111/fwb.14012

Wu, L., Yamamoto, Y., Yamaguchi, S., Minamoto, T., 2022. Spatiotemporal changes in environmental DNA concentrations caused by fish spawning activity. Ecological Indicators 142, 109213. 10.1016/j.ecolind.2022.109213

Xie, S., Watanabe, Y., Saruwatari, T., Masuda, R., Yamashita, Y., Sassa, C., Konishi, Y., 2005. Growth and morphological development of sagittal otoliths of larval and early juvenile *Trachurus japonicus*. Journal of Fish Biology 66, 1704–1719.

Yamamoto, S., Minami, K., Fukaya, K., Takahashi, K., Sawada, H., Murakami, H., Tsuji, S., Hashizume, H., Kubonaga, S., Horiuchi, T., Hongo, M., Nishida, J., Okugawa, Y., Fujiwara, A., Fukuda, M., Hidaka, S., Suzuki, K.W., Miya, M., Araki, H., Yamanaka, H., Maruyama, A., Miyashita, K., Masuda, R., Minamoto, T., Kondoh, M., 2016. Environmental DNA as a ‘snapshot’ of fish distribution: A case study of Japanese Jack Mackerel in Maizuru Bay, Sea of Japan. PLOS One 11, e0149786. 10.1371/journal.pone.0149786

Yoda, M., Mizuta, K., Matsuyama, M., 2006. Induction of ovarian maturetion and ovulation in jack mackerel *Trachurus japonics* by human chorionic gonadotropin. Bull. Fish. Res. Agen 16, 15–18.

