## Supplementary figures and images for "Assessing the potential of eDNA analysis for spawning surveys in marine environments: A case study on Japanese jack mackerel"

### Supplemental figure

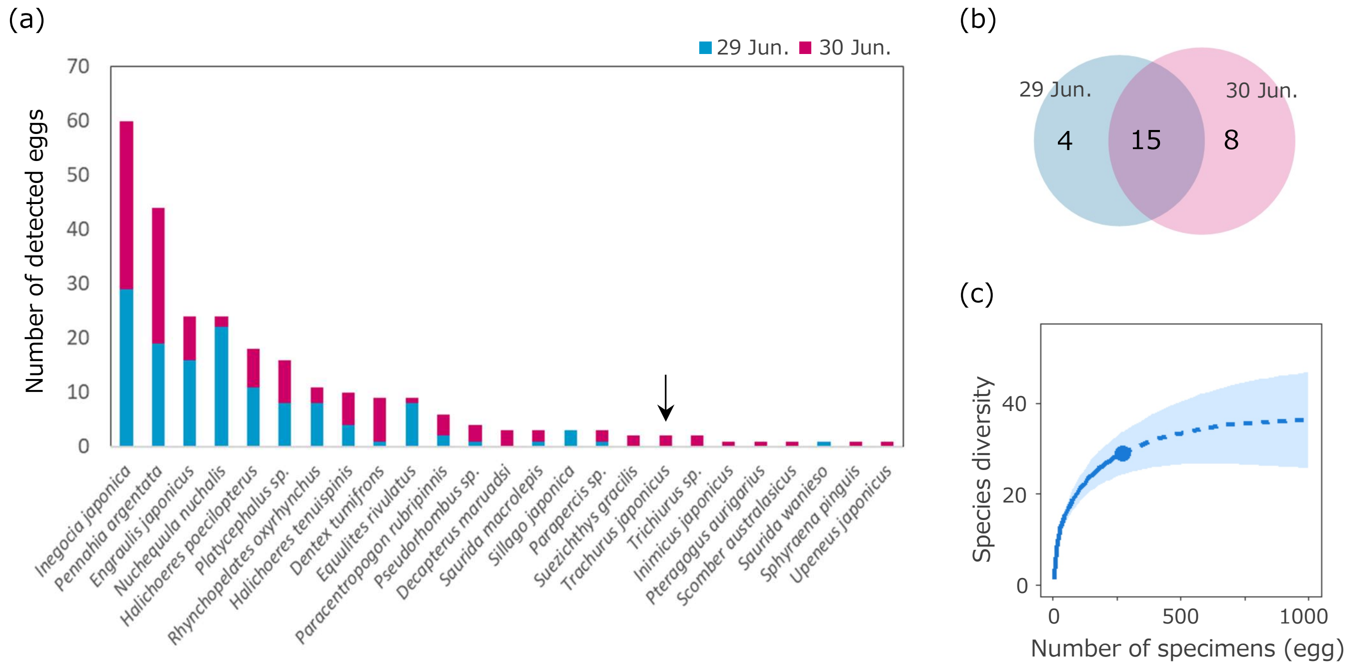
